# Converting Spectral Evoked-to-Background Ratio into Time-Domain Signal-to-Noise Ratio – Validation for High Frequency Oscillations

**DOI:** 10.1101/2025.06.30.662283

**Authors:** G. Fischer, J. Haueisen, D. Baumgarten, M. Kofler

## Abstract

*N* -Interval Fourier Transform Analysis (*N* -FTA) allows for spectral separation of an evoked target signal from uncorrelated background activity. It computes the frequency-dependent evoked-to-background ratio (EBR). The developed method allows for conversion of the spectral EBR into expected values for improvement of signal-to-noise ratio with progressing sweep count. Our study presents the mathematical basis for this conversion along with a validation for simulated and recorded data. The major findings are:

Three factors enter the calculus of the expected SNR: the ratio of durations of the single sweep cycle and the evoked response window, the mean EBR in the spectral target band, and the sweep count. By conversion of all factors to dB, the expected SNR is defined by their sum.
The two fundamental theories governing the improvement of SNR with increasing sweep count, the law of large numbers and the uncertainty principle of signal processing, deliver identical results.
Conversion of EBR to expected SNR was successfully validated by simulated and recorded data and can be applied to all types of evoked data.

**Graphical Abstract:** 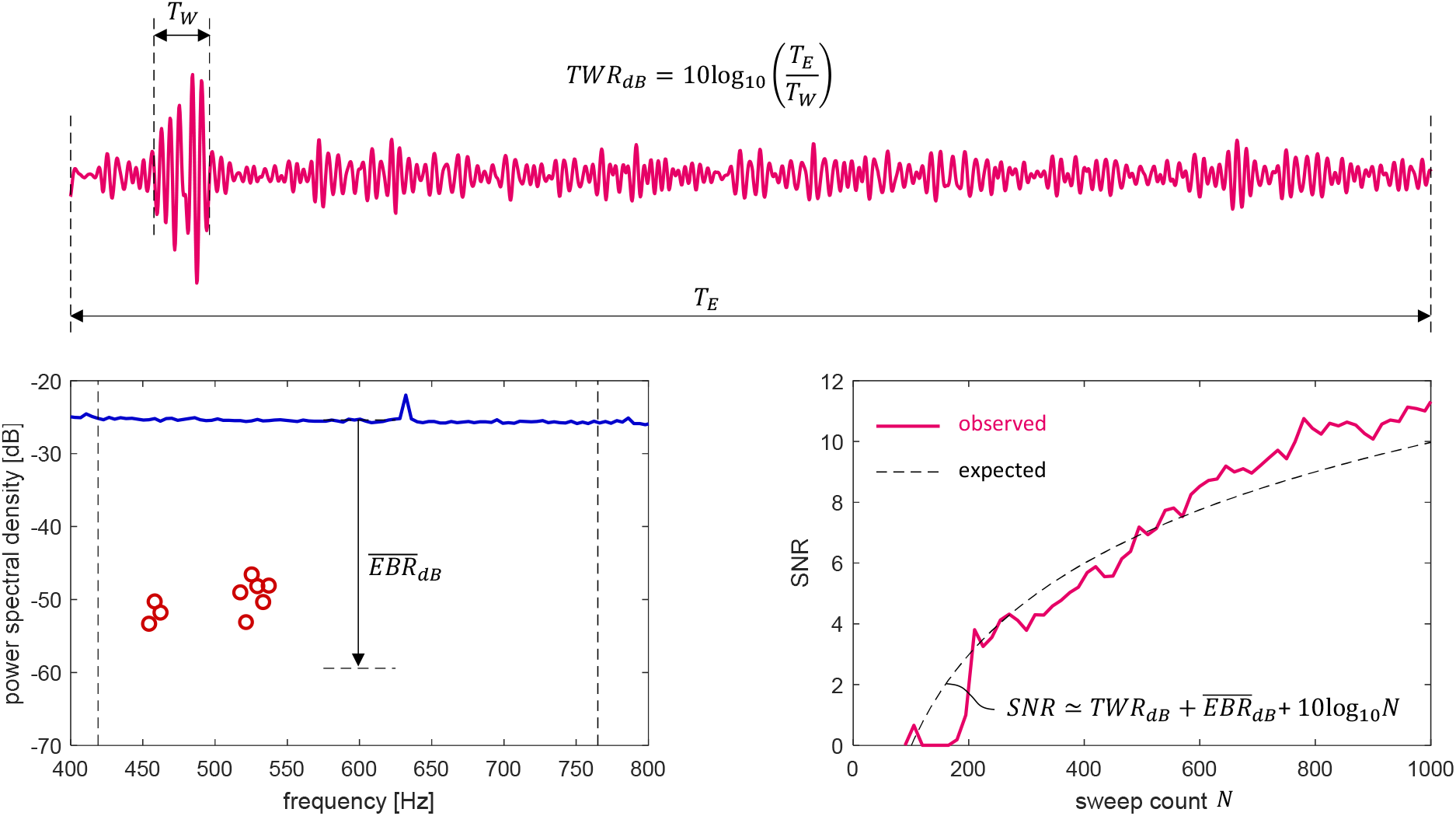

## Background

Trial averaging is used for extracting stimulus-locked somatosensory evoked potentials (SEPs) of a lower amplitude from spontaneous background activity of a larger amplitude [4, 6]. The general mathematical theory of trial averaging is described in [8]. However, signal extraction also involves target band filtering. Therefore, a method is needed that links the spectral properties of both evoked and spontaneous biosignal components with the obtainable response quality in individual recordings.

*N* -Interval Fourier Transform (N-FTA) – a novel spectral separation approach – allows for simultaneous assessment of evoked and background spectral power density (PSD) [1, 2, 3]. Here, the spectral evoked-to-background ratio (EBR) is related to the time-domain signal-to-noise ratio (SNR). This study presents a method that allows for predicting the progress of SNR in individual recordings with increasing sweep count by using the mean target band EBR. Since *N* -FTA reflects the mean PSD levels in some minutes of data, the obtained results of our method reflects the expected values for SNR during a series of sweeps. Theory and data are presented such, that findings are comparable with an actual guideline for recording intraoperative SEPs [7]. The method is validated by a computer simulation and experimental data taken from the original method publication [2].

## Methods Details

### Noise Suppression by Trial Averaging

#### Time domain

As shown in [8], trial averaging reduces the variance in a stochastic signal by division by *N*. When applying target band filtering, the direct current (DC) component is removed. Thus, the variance of the target band filtered response equals its mean power 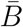. The expected value 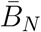 for the noise power at *N* sweeps becomes

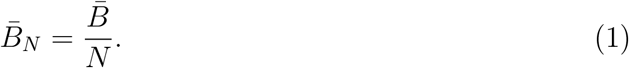

Notably, the relation of variance and sample size *N* is also known as the law of large numbers - a powerful law in statistics [12]. It is independent of the actual statistical distribution. Thus, eq. (1) applies to all kinds of stationary stochastic noise.

#### Frequency domain

The mean power spectral density (PSD) 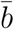 of a target band filtered signal is defined by the fraction of mean noise power 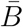 and bandwidth Δ*f*

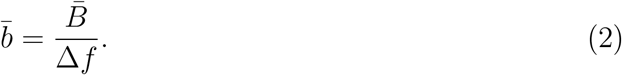

Spectral analysis by *N* -FTA allows for experimental assessment of 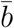. Trial averaging acts like a comb filter allowing all harmonics of *f*_*E*_ to pass while suppressing spectral components in between the harmonics [8]. The trial averaging transfer function *H*_*N*_ (*f*) is defined by

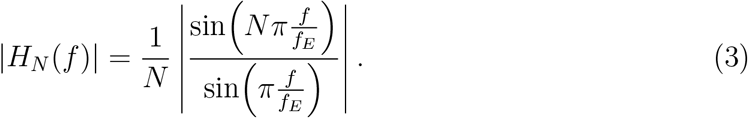

As can be taken from Figure 1, trial averaging generates a periodic transfer function containing distinct peaks at the harmonics. The remaining noise 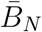 after N averages is obtained from the integral over the power at the output of the trial average filter

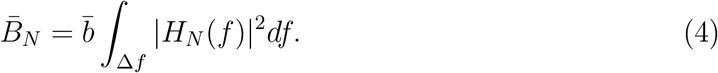

**Figure 1:**
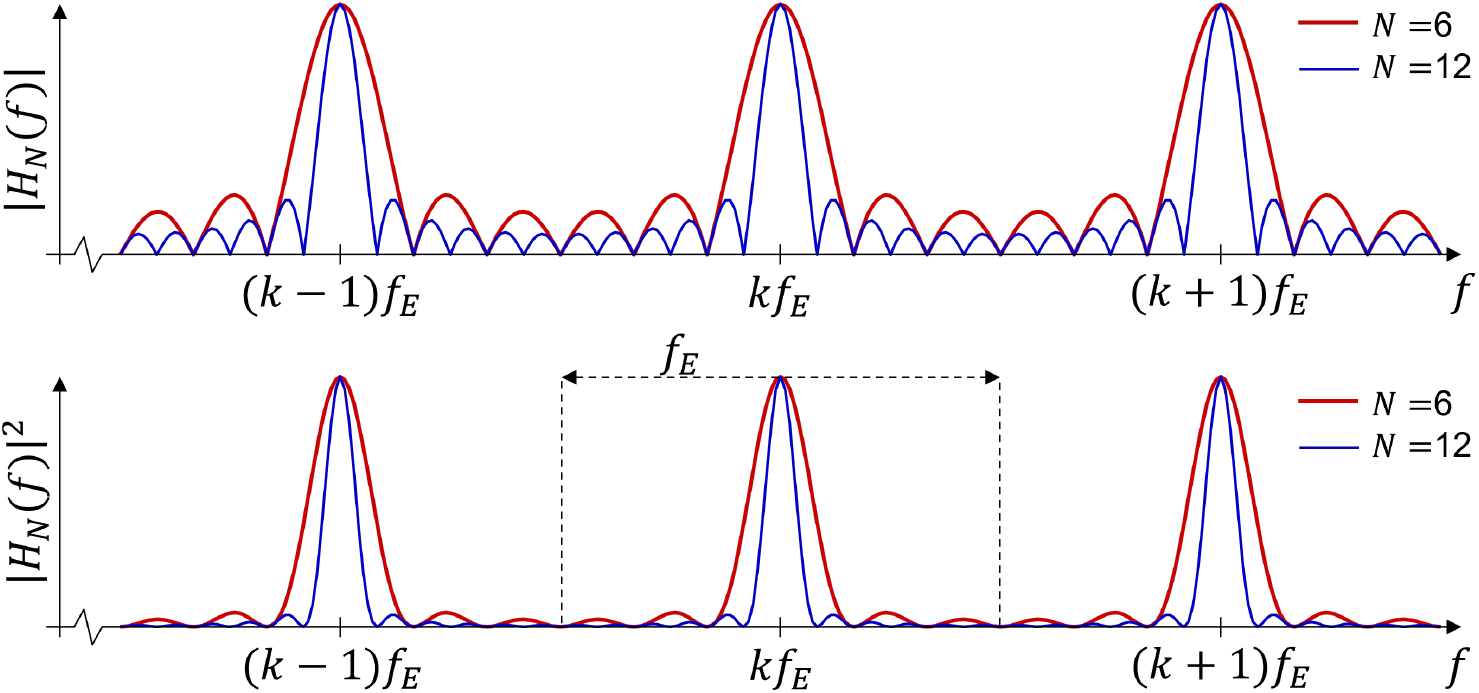
*Top:* Magnitude of the periodic transfer function |*H*_*N*_ (*f*) | obtained for trial averaging. A sweep count *N* of six and twelve was chosen for illustration. Trial averaging acts as comb filter with peaks at integer multiples of the stimulation rate *f*_*E*_. Between these peaks the signal gets attenuated. With increasing sweep count peaks get narrowed and a better attenuation is obtained. *Bottom:* The square of the transfer function governs the power transmission of a stochastic signal. The dashed lines mark one cycle as chosen for the auxiliary calculation (see text).

Here, the notation chosen in eq. (4) indicates that integration is performed over the pass-band of the target-filter (i.e., from the lower to the upper corner frequency yielding the bandwidth Δ*f*). For solving the integral in eq. (4) it is convenient considering integration over a single cycle *f*_*E*_ of the periodic spectrum

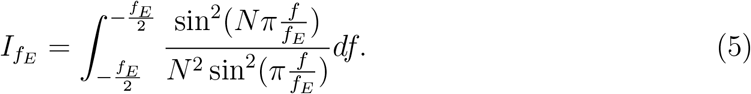

The solution of the integral over a single cycle is solved by an auxiliary calculation at the end of this section yielding

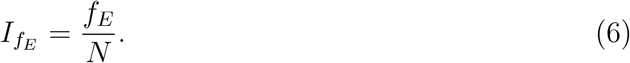

By observing that the target band Δ*f* contains Δ*f/f*_*E*_ periodic cycles we obtain for the integral in eq. (4)

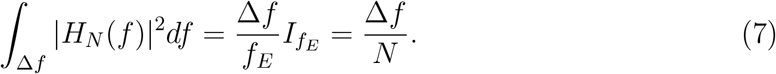

Substituting (7) back into (4) we obtain for noise suppression after *N* sweeps

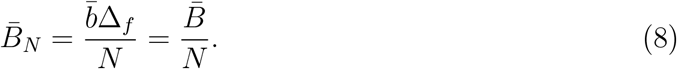

#### Interpretation

Consistently, time domain analysis by eq. (1) and frequency domain analysis by eq. (8) provide identical results. Most importantly, this double analysis provides a useful observation. In the time domain the law of large numbers explains the mechanism of noise suppression. In the frequency domain, increasing *N* decreases the width of the combs in *H*_*N*_ (decreasing uncertainty in frequency resolution). Thus, our analysis links the law of large numbers with the uncertainty principle of signal processing [9] - two general theories, both delivering identical results.

#### Target-Signal Power

In a stationary experimental setting the evoked response can be modeled as a finite periodic signal. Since its harmonics concur with the peaks of the combs, trial averaging does not alter the signal. Thus, the average power 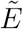 of the evoked signal is obtained from the mean evoked PSD *ē* within the target band Δ*f*

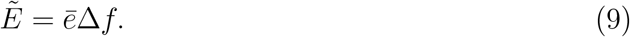

Importantly, 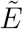 is the average of evoked power taken over the entire evoked cycle *T*_*E*_ (see Figure 2). The power of evoked HFOs however, is distributed unevenly within a cycle *T*_*E*_ and almost entirely contained in the narrow evoked response window *T*_*W*_. The mean power *Ē* within the response window *T*_*W*_ is obtained from

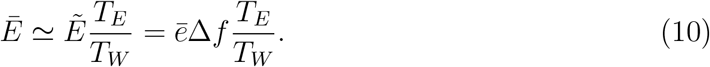

**Figure 2:**
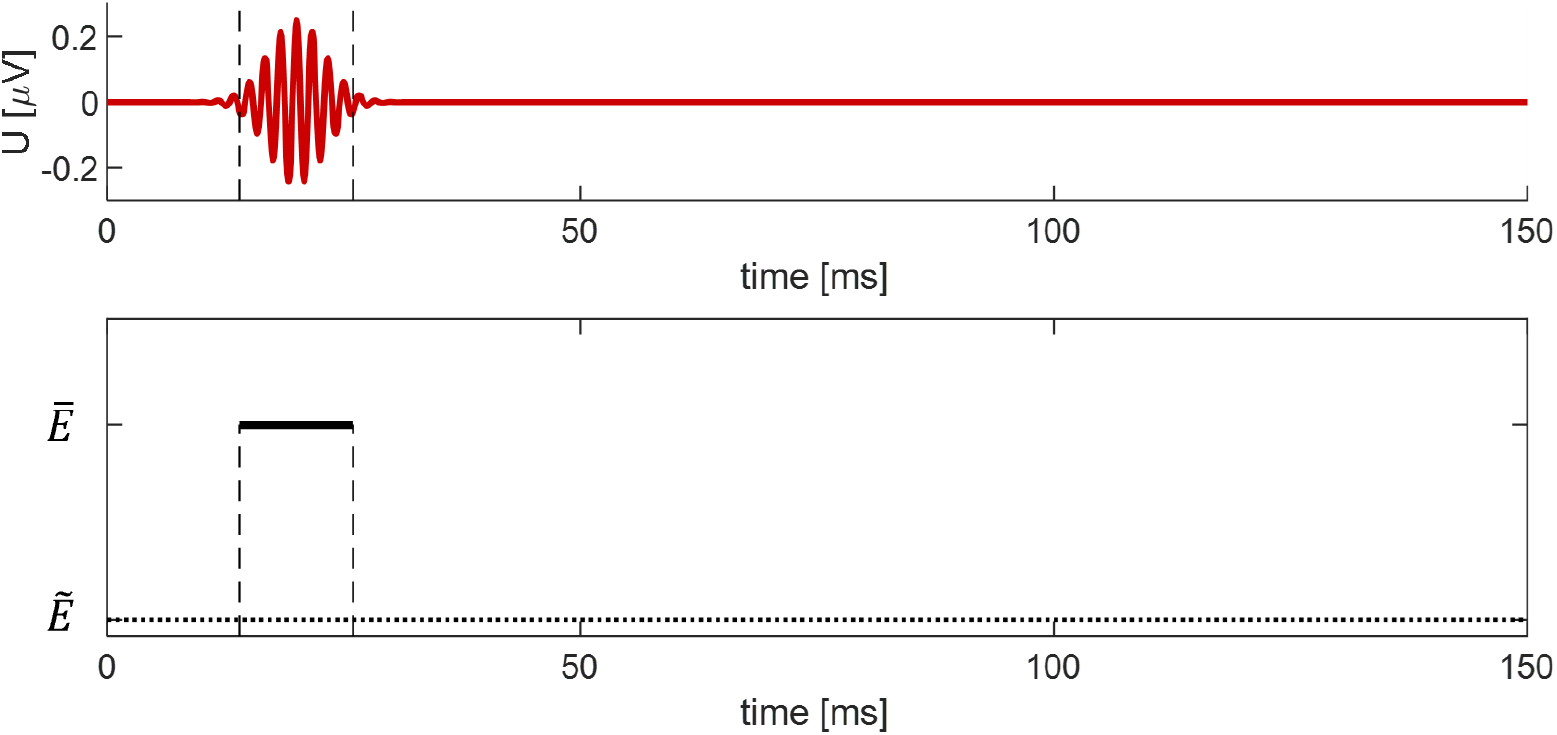
*Top:* Schematic representation an HFO of width *T*_*W*_ (dashed line) within a cycle *T*_*E*_ of 150 ms duration. *Bottom:* 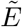 is the mean HFO-power within *T*_*E*_, while *Ē* is its mean power within *T*_*W*_.

In an experimental setting, the choice of *T*_*W*_ involves some uncertainty. Therefore, equation 10 should be considered as an approximation.

#### Signal-to-Noise Ratio

Taking advantage of equations (8) and (10) the expected value for the target band SNR is now defined by

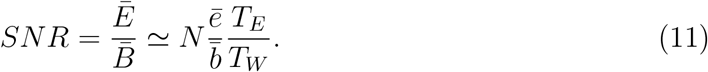

Guidelines request the conversion of SNR to dB [6, 7]

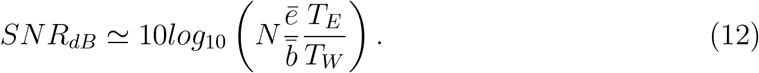

Notably, sweep count *N* enters SNR linearly which corresponds to a logarithmic curve in a dB scale. Equation (12) can be rewritten to express SNR as a sum of three terms containing the time window ratio *T*_*E*_*/T*_*W*_, the mean evoked-to-background ration 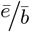, and sweep count *N*

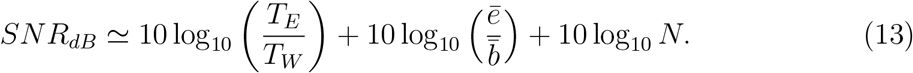

#### Interpretation

The time window ratio (first term in eq. (13)) is always positive and shifts the SNR towards higher values. The mean EBR is typically negative (high level background activity). Thus, the second term in eq. (13) shifts the SNR towards lower values. The actual improvement of SNR with increasing sweep count depends only on *N* (third term in eq. (13)) being, therefore, independent from individual physiological parameters.

#### Auxiliary Calculation – Power Transfer in a Single Comb Cycle

Power transfer in a single comb cycle (Figure 1, bottom panel) is obtained from the solution of the integral in eq. (5). By substituting 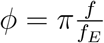 this integral can be rewritten

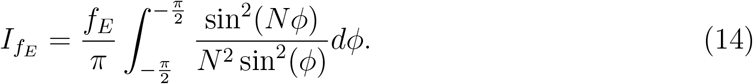

For solving the integral in eq. (14) a first approximation is made by replacing the denominator sin *ϕ* by *ϕ*

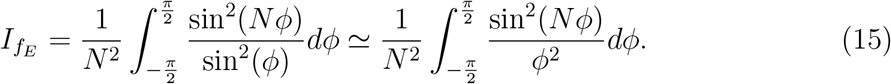

As can be taken from Figure 3, this approximation allows for an accurate estimation of the integral for large *N*. The sinc-squared function in (15) can be solved using integration by parts (taking 1*/ϕ*^2^ as the part to integrate)

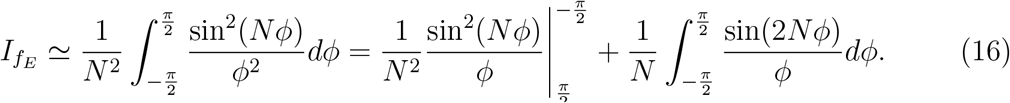

**Figure 3:**
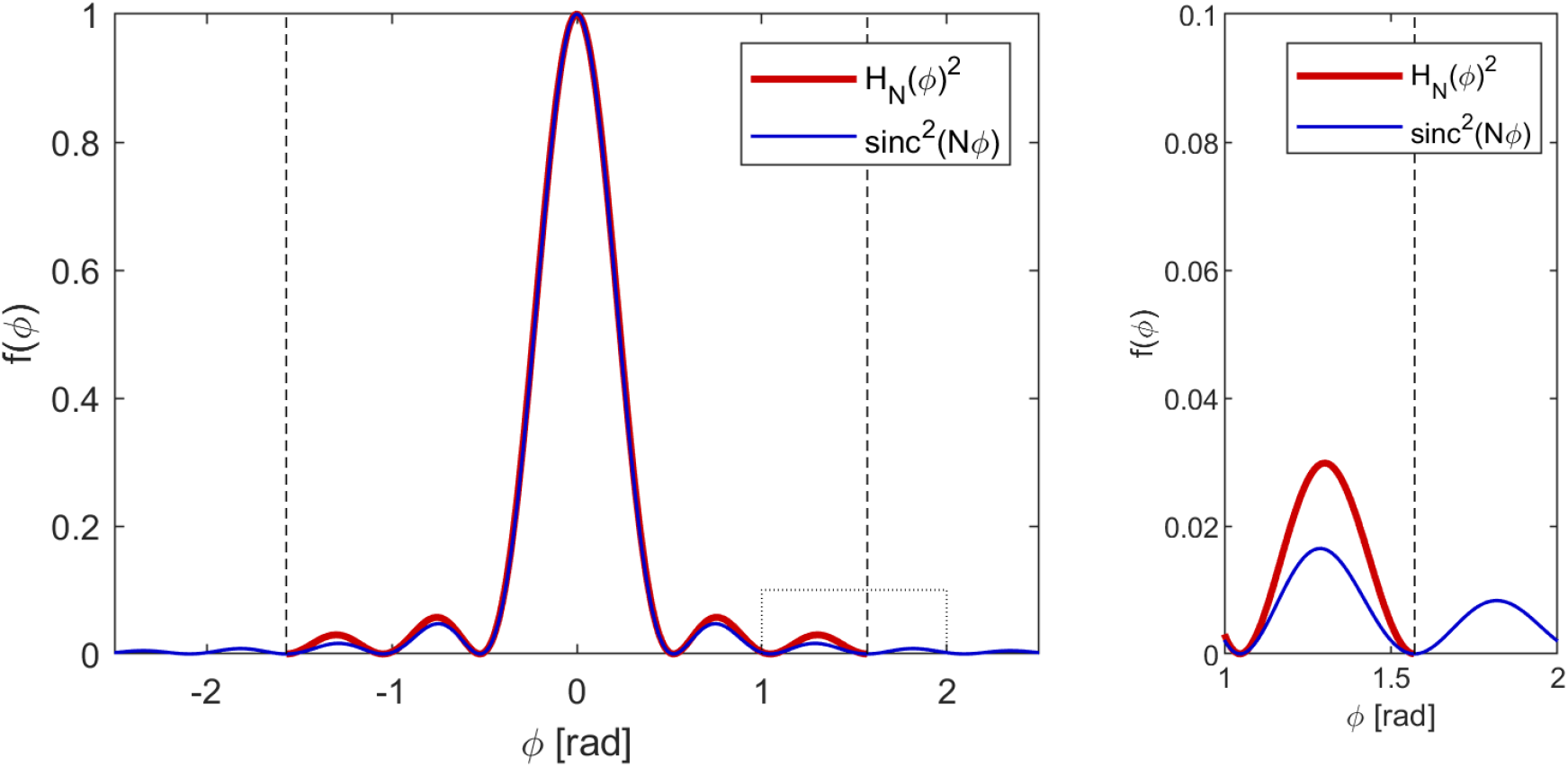
*Left:* The square of the transfer function *H*_*N*_ (*ϕ*) (eq. (14), red) contains sin(*ϕ*) in the denominator. Replacement of this term by *ϕ* yields the square of a sinc-function (eq. (15), blue). Near the origin both functions are in good agreement while near 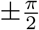 (vertical dashed lines) small but visible deviations can be observed. *N* = 6 was chosen for illustration. *Right:* Magnification of the dotted area in the left panel. The square of the sinc-function is of smaller values compared to the square of the transfer function but extends over a broader interval. These two deviations compensate each other when computing the integral.

The first term on the right hand side of eq. (16) approaches zero for large *N* due to the division by *N* ^2^. Therefore, this term can be neglected for large *N*. For solving the integral on the right hand side of eq. (16) the substitution *u* = 2*Nϕ* is applied

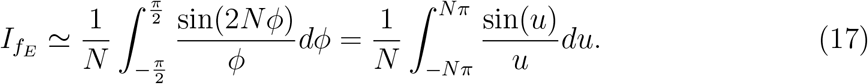

For large *N* integration is now carried out over multiple oscillations of sinc(*u*). As a second approximated integration is made from *−∞* to *∞*. It is a known property of the sinc-function that its infinite integral yields *π*. We obtain

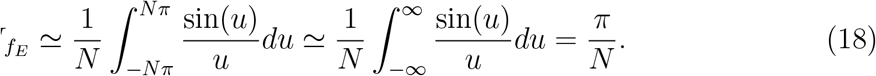

The calculus performed above suggests that 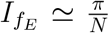 provides an accurate estimate for large *N*. We investigated also some low values of *N*, which allow for straight forward integration. For *N* = 1 the term within the integral of eq. (14) reduces to one. Thus, we obtain 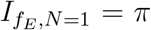 which exactly agrees with eq. (18). For *I* = 2 integration can be simplified by using sin(2*ϕ*) = 2 sin *ϕ* cos *ϕ* eq. (15). We obtain by integration by parts

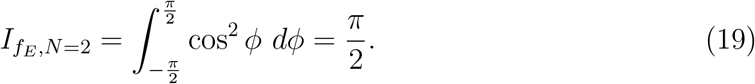

Again, this agrees exactly with eq. (18). Furthermore, when performing numerical integration for *N* = 3, …, 20, it turned out that results matched the prediction of eq. (18) accurately. The magnitude of the relative error (numerical integration vs.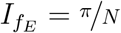) was always below 10^−15^ and, thus, at the resolution of standard 32-bit floating point numbers.

Concluding, our analysis confirms that eq. (18) provides not only an estimation but the exact value for the integral. Figure 3 provides an explanation for this observation. The first approximation (substitution of sin(*ϕ*) by *ϕ*) slightly underestimates 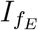 (narrow blue trace in the figure). The second approximation (integration over ±∞ instead of 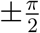) slightly overestimates 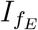 (extension of integration interval). Our investigations showed that these two approximations exactly compensate for each other.

## Method Validation

This section aims to validate eq. (13) as the key result of our analysis by means of simulated and recorded data. Validation requires reliable estimates of the actual observed SNR. For obtaining these estimates, a first time window *T*_*W*_ (response window) is selected containing a superposition of evoked and spontaneous power *E* and *B*, respectively (Figure 4). Here, power is the mean square of the amplitude within the window *T*_*W*_. Importantly, only the sum Σ = *E* + *B* of these two variables is accessible.

**Figure 4:**
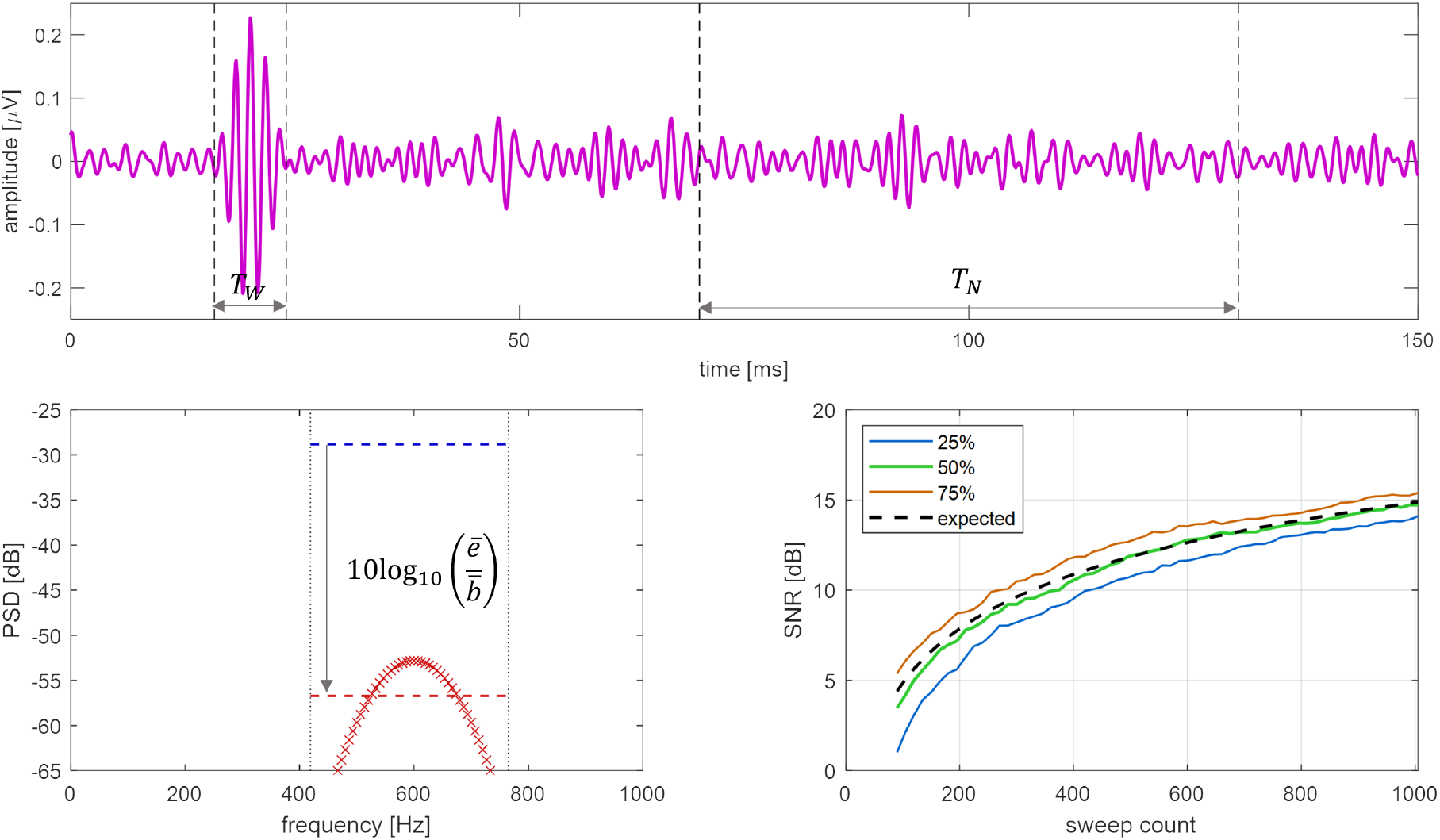
*Top:* Example for an extracted response (*N* = 1005, zero-phase IIR filter). The time intervals applied for computing the observed SNR (13.4 dB for the shown example) are indicated (vertical dashed lines). *Bottom Left:* The PSD for the Morlet wavelet is depicted by red markers. Within the target band (vertical dotted lines) the mean evoked PSD (red dashed line) and mean background PSD (blue dashed line) are shown. The negative EBR is indicated by a downwards arrow. *Bottom Right:* The quartiles for the observed SNR (zero-phase IIR filter) are shown together with the expected SNR.

As outlined in [2], the background power level can be estimated from a long segment at the end of the evoked cycle. Thus, a second time window *T*_*N*_ (noise window) is applied to obtain *B* (Figure 4). An estimate for the SNR at *N* sweeps is obtained from

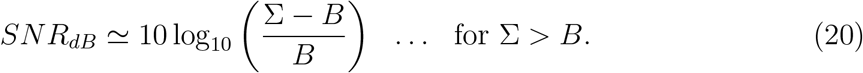

At high noise levels (e.g., at a low sweep count *N*) *B* may exceed Σ due to statistical variations making results meaningless. Hence, validation should be performed at sufficiently large *N*. Furthermore, also SNR estimates containing negative dB values are of limited use, since such data are dominated by noise. Therefore, the implemented code processes only positive SNR values. A minimal sweep count of *N* = 90 was selected and SNR was analyzed at steps Δ*N* = 15 up to *N* = 1005.

Two different types of HFO target band filters were used in the validation process. Firstly, the zero-phase bandpass, which was originally proposed in [5] and also used in [2] was tested (419 to 765 Hz bandwidth). Secondly, two Ricker wavelets ([11], 585 Hz center frequency) were applied in series. Here, a bandwidth of 418 to 769 Hz was obtained, allowing for direct comparison of both settings.

### Simulated Data

Simulations allow for accurate testing since parameters for both the evoked response and for random background activity can be predefined and verified. A Morlet wavelet (see original method publication [2]) was chosen as a surrogate evoked HFO. Briefly, a Morlet wavelet is an oscillation of a predefined frequency *f*_*e*_ and a bell-shaped envelope of amplitude *V*_*e*_ with a standard deviation (time constant) *τ*_*e*_. White noise of standard deviation *V*_*N*_ was added as a surrogate of background activity. Data were sampled at the rate *f*_*S*_. The selected parameters are listed in Table 1. Notably, background noise exceeds the evoked signal considerably.

**Table 1:** Parameters for simulated HFOs.

|  |  |
| --- | --- |
| chosen parameters |  |
| evoked amplitude $V_e$ | 0.25 μV |
| evoked frequency $f_e$ | 600 Hz |
| evoked envelope standard deviation $\tau_e$ | 2 ms |
| response duration $T_W$ | $4 \times \tau_e = 8$ ms |
| evoked cycle duration $T_E$ | 150 ms |
| noise window $T_N$ | 60 ms |
| white noise standard deviation $V_N$ | 2.5 μV |
| sample rate $f_S$ | 9.6 kHz |
| computed ratios |  |
| time window ratio | 12.7 dB |
| mean EBR | -27.9 dB |
| expected SNR |  |
| $N = 1$ (SNR without trial averaging) | -15.1 dB |
| $N = 100$ | 4.9 dB |
| $N = 1000$ | 14.9 dB |

In total 100 series each containing 1005 sweeps were simulated. The top panel of Figure 4 shows a simulated result. The time window *T*_*W*_ was centered at the highest peak of the Morlet wavelet. The time window *T*_*N*_ was placed such that its right end is 20 ms ahead of the end of the evoked interval.

The evoked PSD was computed by applying an FFT to the Morlet wavelet. Within the spectral target band a mean evoked PSD of *−* 56.7 dB was obtained (Figure 4, left bottom panel). White noise PSD was 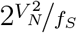 corresponding to *−* 28.8 dB. By inserting these values into eq. (13) the expected SNR values in Table 1 and in the bottom right panel of Figure 4 were obtained.

The observed SNR was calculated for the 100 simulated sweeps (eq. (20)) and its quartiles were obtained. When comparing the median of the observed SNR (zero-phase IIR filter) with the expected SNR a correlation coefficient of 0.999 and a root-mean-square deviation of 0.3 dB were obtained. The interquartile range in the observed SNR was about 2 dB.

When performing the same test using the Ricker-wavelet bandpass, a similar result was obtained. Here observed SNR was shifted by about 0.4 dB towards lower values. Again, a correlation coefficient of 0.999 was obtained, but root-mean-square deviation increased to 0.7 dB. The interquartile range was again 2 dB. This moderate reduction of SNR was potentially caused by the smoother transition of the Ricker wavelet transfer function to its stop bands (as compared to the zero-phase IIR filter). This allowed for slightly more noise transfer in the filters stop-bands.

Concluding, both investigated bandpass filters allowed for a successful validation of the method by computer simulation. The zero-phase IIR filter produced slightly better results.

### Recorded Data

The evoked spectral PSD of HFOs is at relatively low levels close to the theoretical limit of resolution of *N* -FTA [2]. Therefore, a recording containing high evoked amplitudes was considered (subject B in our original method publication [2], stimulation rate *f*_*E*_=3.95 Hz). The response window *T*_*W*_ was placed at 15 to 25 ms. For the noise window *T*_*N*_ a width of 60 ms was chosen again and also here, the window was placed 20 ms ahead of the end of the evoked cycle. *N* -FTA delivered a mean EBR of −34 dB (see online Graphical Abstract, left bottom panel).

The tested filters were identical as for the simulated data. For both settings the observed SNR was in good agreement with the expected SNR (Figure 5). A quantitative comparison of the expected SNR with the observed SNR (zero-phase IIR bandpass) revealed a correlation coefficient of 0.982 and a root-mean-square deviation of 1.0 dB. Notably, this deviation obtained for a single recording is well within the interquartile range observed in the simulations. When comparing the results obtained from the two tested filters, the observed SNR displayed good agreement (correlation coefficient 0.999; root-mean-square deviation 0.2 dB).

**Figure 5:**
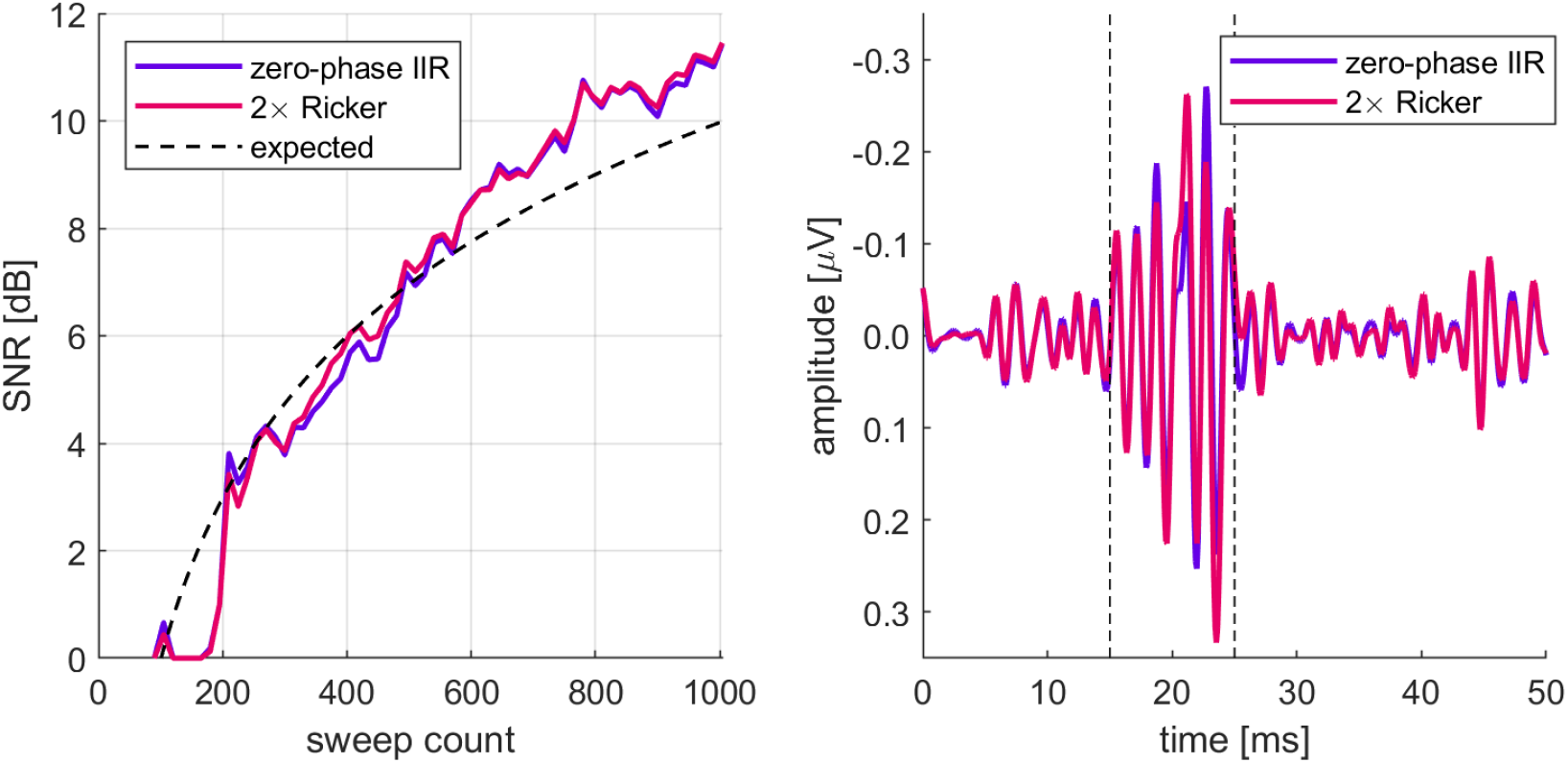
*Left:* Observed improvement of SNR for the two investigated bandpass filters (full lines) and expected SNR obtained from spectral analysis (dashed line). *Right:* Evoked HFO responses obtained by the two filters. The vertical dashed lines mark the response interval applied for computing the expected SNR.

Also, the evoked HFO-responses obtained from the two filters were in good agreement (correlation coefficient 0.95 within the evoked response window *T*_*W*_). In between 20 to 25 ms a small visible displacement of the traces was observed. This was caused by the differences in attenuation of the tested filters in the high-γ-band (zero-phase IIR filter has better attenuation than the Ricker wavelet). As shown in [1] cortical responses to median nerve stimulation contain significant activity in the high-γ-band. The zero-phase IIR bandpass provides a better suppression of these high-γ-band components.

### Individual variations

Within the HFO-band, recorded data display a remarkable individual variation in mean EBR. In the data investigated in [2], the average value was near *−*38 dB. From the data presented in [10], it appears reasonable to choose at least three times the cycle length of an HFO for *T*_*W*_. This yields approximately 5 ms, while the evoked interval *T*_*E*_ equals some hundred ms. This increases SNR by about 15 dB. Thus, the mean target band SNR at *N* = 1 may be near −23 dB. This allows for estimating that roughly 2000 sweeps (equal to an increase of 33 dB) are required for extracting an HFO response at an SNR of 10 dB. Notably, when assuming that the mean EBR displays a variation of *±* 5 dB (this corresponds approximately to doubling/halving HFO-amplitude) the sweep count required for obtaining 10 dB may vary between roughly 600 and 6000 repetitions.

Concluding, in a data set providing high-quality spectral separation, a successful validation was performed. A consideration of individual variation provides reasonable estimates for required sweep counts being in agreement with observations from human experiments. Our method is useful for the online estimation of the individually required number of trials in any averaging procedure of evoked potentials of fields.

## Ethics Statements

The experimental data was taken from a study approved by the institutional *Research Committee for Scientific Ethical Questions* (RCSEQ 2632/19). Informed consent was obtained from all subjects.

## CRediT Author Statement

Gerald Fischer: Methodology, Software, Writing, Original draft preparation. Jens Haueisen: Reviewing and Editing. Daniel Baumgarten: Resources, Reviewing and Editing. Markus Kofler: Investigation, Writing, Reviewing and Editing.

## Acknowledgments

GF wants to thank his former colleague Michael Seger for stimulating discussions on bandpass filters. This work received no funding.

## Supplementary Material and Additional Information

### Supplementary Material

Recorded data and the MATLAB code are available for download (see Specifications Table).

